# Electrophysiological evidence for impaired auditory sensory memory in Cystinosis despite typical sensory processing: An MMN investigation

**DOI:** 10.1101/747642

**Authors:** Ana A. Francisco, John J. Foxe, Douwe J. Horsthuis, Sophie Molholm

## Abstract

Cystinosis, a genetic rare disease characterized by cystine accumulation and crystallization, results in significant damage in a multitude of tissues and organs, such as the kidney, thyroid, eye, and brain. While Cystinosis’ impact on brain function is relatively mild compared to its effects on other organs, the increased lifespan of this population and thus potential for productive societal contributions have led to increased interest on the effects on brain function. Nevertheless, and despite some evidence of structural brain differences, the neural impact of the mutation is still not well characterized.

Here, using a passive duration oddball paradigm (with different stimulus onset asynchronies (SOAs), representing different levels of demand on memory) and high-density electrophysiology, we tested basic auditory processing in a group of 22 children and adolescents diagnosed with Cystinosis (age range: 6-17 years old) and in neurotypical age-matched controls (N=24). We examined whether the N1 and mismatch negativity (MMN) significantly differed between the groups and if those neural measures correlated with verbal and non-verbal IQ. Individuals diagnosed with Cystinosis presented *similar* N1 responses to their age-matched peers, indicating typical basic auditory processing in this population. However, whereas both groups showed similar MMN responses for the shortest (450ms) SOA, suggesting intact change detection and sensory memory, individuals diagnosed with Cystinosis presented clearly reduced responses for the longer (900ms and 1800ms) SOAs. This could indicate reduced duration auditory sensory memory traces, and thus sensory memory impairment, in children and adolescents diagnosed with Cystinosis. Future work addressing other aspects of sensory and working memory is needed to understand the underlying bases of the differences described here, and their implication for higher order processing.

## Introduction

Cystinosis, caused by bi-allelic mutations in the 17p13.2-located CTNS gene (Town et al., 1998), is an autosomal recessive disorder with an incidence of approximately one in 100,000 to 200,000 live births (Gahl, Thoene, & Schneider, 2009). Though over 100 mutations have been identified, the most common is a 57-kb deletion (Levtchenko, van den Heuvel, Emma, & Antignac, 2014; Shotelersuk et al., 1998). CTNS encodes cystinosin, a lysosomal cystine-proton co-transporter. Its mutation results in excessive cellular cystine storage (Gahl, Bashan, Tietze, Bernardini, & Schulman, 1982; Jonas, Greene, Smith, & Schneider, 1982), which appears to cascade into deregulation of endocytosis and cell signaling processes (Ivanova, De Leo, De Matteis, & Levtchenko, 2014). Moreover, intralysosomal cystine crystallizes, triggering significant damage in a multitude of tissues and organs (Gahl & Kaiser-Kupfer, 1987).

The first manifestations of the disease emerge at around six months of age (Gahl, 1986), with typical development being described until then. Amid other possible complications, CTNS mutations often result in end-stage renal disease, hypothyroidism, and retinopathy (Vogel et al., 1990), at least in Infantile Nephropathic Cystinosis, the classic and more prevalent form of the disorder (Schneider, Katz, & Melles, 1990), and the one addressed in the present study. Despite the undoubtedly multi-systemic nature of the disease (Elmonem et al., 2016), effectively treating the associated renal complications was the obvious focus until approximately 20 years ago. The emergence of renal replacement therapy and the development of cysteamine, a cystine-depleting agent which slows the progression of renal failure and protects extra-renal organs (van Rijssel, Knuijt, Veys, Levtchenko, & Janssen, 2019), greatly increased life expectancy in this population (now above 50 years (Ivanova et al., 2014)), and allowed for a more prominent focus on the characterization of other aspects of the disease, such as the neurological, cognitive, and behavioral sequelae.

Human studies have since shown abnormally high levels of cystine in various brain regions (Levine & Paparo, 1982; Theodoropoulos, Krasnewich, Kaiser-Kupfer, & Gahl, 1993; Vogel et al., 1990), and long-term adverse effects of Cystinosis on the central nervous system (Niemiec, Ballantyne, & Trauner, 2012). Furthermore, different neurological findings have been described, which include subcortical and cortical atrophy, Chiari I malformation, white matter abnormalities, and atypical electrophysiological (EEG) activity (Bava et al., 2010; Cochat, Drachman, Gagnadoux, Pariente, & Broyer, 1986; Ehrich, Stoeppler, Offner, & Brodehl, 1979; Fink et al., 1989; Rao, Hesselink, & Trauner, 2015). Cognitively, individuals diagnosed with Cystinosis often present intelligence quotients (IQ) in the typical range, but lower IQs have also been reported (Aly, Makar, El Bakri, & Soliman, 2014; Ulmer et al., 2009). A differential between non-verbal and verbal indices is consistently reported in this population, with the former being significantly lower than the latter (Frankel & Trauner, 2019; Spilkin, Ballantyne, Babchuck, & Trauner, 2007; Ulmer et al., 2009). This pattern appears to emerge early in development and to persist throughout the lifespan (Scarvie, Ballantyne, & Trauner, 1996; Trauner, Chase, Scheller, Katz, & Schneider, 1988), regardless of age at treatment onset (Viltz & Trauner, 2013). Significant difficulties are observed in visual-motor, visual-spatial and visual memory skills, as well as executive function related abilities (Aly et al., 2014; Ballantyne, Spilkin, & Trauner, 2013; Besouw et al., 2010; Sathappan & Trauner, 2019; Viltz & Trauner, 2013), which may particularly hinder academic skills (Ballantyne, Scarvie, & Trauner, 1997). Motor deficits and fine motor incoordination have also been described (Trauner, Spilkin, Williams, & Babchuck, 2007; Trauner et al., 2010; Ulmer et al., 2009). Some of these difficulties seem to be likewise present in unaffected heterozygous carriers of the cystinosin gene mutation (Sathappan & Trauner, 2019).

Despite compelling evidence that CTNS mutations are associated with structural brain differences and cognitive impairments, Cystinosis’ impact on brain activity is still not well understood. High-density EEG, a non-invasive method that provides information at the millisecond scale, allows one to directly measure functional brain activity and thus reliably assess the integrity of neural function. The sparseness of studies in which EEG has been used to assess functional brain activity in Cystinosis is, thus, quite surprising. One case study looked at auditory and somatosensory evoked potentials in an adult female. Though no methodological details or specific result were included, typical neural activity was reported (Müller, Baumeier, Ringelstein, & Husstedt, 2008). A conference paper reported an enhanced auditory P2 for 14 individuals diagnosed with Cystinosis (age range: 6 to 52 years old), during a spatial localization task (Čeponiene et al., 2008). A more recent case study tested visual processing in two children with Cystinosis before and after kidney transplantation. Before transplantation (and during dialysis treatment), both children showed delayed and decreased early visual-evoked responses, when compared to their age□matched peers. Remarkably, both amplitude and latency measures normalized upon retest, two years after transplant (Ethier, Lippé, Mérouani, Lassonde, & Saint-Amour, 2012). In spite of the paucity of studies and the very small number of individuals tested to date, EEG measurements seem nonetheless to be sensitive to neuropathology in Cystinosis. Importantly, EEG and event-related potentials (ERPs) may be leveraged as outcome measures to assess the impact of treatment on brain function vis-à-vis neurophysiological integrity.

Therefore, here, to gain insight into potentially impaired neural function, we used high-density EEG to assay basic sensory processing in Cystinosis. We focused on early auditory sensory processing (the N1) and sensory memory (the mismatch negativity, MMN). The auditory N1 is the first prominent negative auditory-evoked potential (Näätänen & Picton, 1987), and reflects neural activity generated in and around primary auditory cortex (Giard et al., 1994; Leavitt, Molholm, Ritter, Shpaner, & Foxe, 2007). The MMN, in turn, operating at the sensory memory level, occurs when a repeating stimulus (the standard) in an auditory stream is replaced by a deviant stimulus: Regular aspects of consecutively presented standards form a memory trace; violation of those regularities by a deviant induces the MMN (Näätänen & Winkler, 1999). Occurring 100 to 200 ms following the deviant event, the MMN is thought to reflect the neural processes underlying detection of a pattern violation and updating of a representation of a regularity in the auditory environment (Näätänen & Alho, 1995; Ritter, Deacon, Gomes, Javitt, & Vaughan, 1995; Ritter, Sussman, Molholm, & Foxe, 2002). To impose different levels of demand on the sensory memory system, the rate of presentation was parametrically varied. Additionally, we were interested in understanding how these neural measures related to cognitive function in Cystinosis. To this end and given the idiosyncratic pattern of IQ scores in Cystinosis, we queried the relationship between N1 and MMN and verbal and non-verbal IQ.

## Materials and methods

### Participants

Twenty-five participants diagnosed with Cystinosis (age range: 6-17 years old; M = 11.08; SD = 2.55) and twenty-eight neurotypical controls (NT) (age range: 6-17 years old; M = 11.42; SD = 3.38) were initially recruited. Exclusionary criteria for the NT group included hearing problems, developmental and/or educational difficulties or delays, and neurological problems. Exclusionary criteria for the Cystinosis group included hearing difficulties and current neurological problems. Participants passed a hearing test (below 25dB HL for 500, 1000, 2000, 4000Hz) performed on both ears using a Beltone Audiometer (Model 112). Four individuals diagnosed with Cystinosis were tested at an off-site location and, therefore, no hearing test was conducted. For these individuals, parents reported normal hearing and no history of hearing problems.

Four neurotypical controls presented high-average or superior verbal IQ scores, but borderline non-verbal IQ scores. Exclusion of these individuals seemed to bear no impact either on the EEG grand-averages or on the statistical results. Given the inconsistencies across verbal and nonverbal IQ indices, these four individuals were, nevertheless, excluded from the final sample reported here. Due to illness on the scheduled day of testing, three individuals with Cystinosis were unable to perform the EEG tasks. Because those participants had traveled from out of town and, thus, could not be rescheduled, they were also excluded from the final sample.

All participants signed an informed consent approved by the Institutional Review Board of the Albert Einstein College of Medicine. Participants were monetarily compensated for their time. All aspects of the research conformed to the tenets of the Declaration of Helsinki.

### Experimental Procedure and Stimuli

Testing occurred over a 2-day period and included a cognitive testing session (focused on measures of intelligence: Wechsler Abbreviated Scale of Intelligence (WASI-II); Wechsler, 1999) and an EEG recording session. The EEG paradigm reported here focused on auditory processing, utilizing a traditional duration-MMN oddball paradigm (Brima et al., 2019). Participants sat in a sound- and electrically-shielded booth (Industrial Acoustics Company Inc, Bronx, NY) and watched a muted movie of their choice on a laptop (Dell Latitude E6430 ATG or E5420M) while passively listening to regularly (85%) occurring standard tones interspersed with infrequently occurring deviant tones (15%). These tones had a frequency of 1000 Hz with a rise and fall time of 10 ms, and were presented at an intensity of 75dB SPL using a pair of Etymotic insert earphones (Etymotic Research, Inc., Elk Grove Village, IL, USA). Standard tones had a duration of 100 ms while deviant tones were 180 ms in duration. These tones were presented in a random oddball configuration (excepting that at least two standards preceded a deviant) to yield an MMN. In three blocked conditions, the stimulus onset asynchrony (SOA) was either 450 ms, 900 ms or 1800 ms. Each SOA was presented in blocks of 4 minutes long, composed of 500, 250 or 125 trials respectively. Participants were presented with 14 blocks (2*450 ms, 4*900 ms and 8*1800 ms), resulting in a possible 1000 trials (and 150 deviants) per SOA. For each of the SOA conditions, the presence and magnitude of the MMN was measured by comparing the event-related potentials (ERPs) to the standard and deviant stimuli. For the four individuals diagnosed with Cystinosis whose data were collected off-site, during a family meeting, the equipment was the same used for the remainder of the subjects, but cognitive and EEG data collections were carried out in two regular rooms. An attempt was made to keep the lighting and sound conditions identical to the ones experienced in the lab setting.

### Data acquisition and analysis

EEG data were acquired continuously at a sampling rate of 512 Hz from 71 locations using 64 scalp electrodes mounted on an elastic cap and seven external channels (mastoids, temples, and nasion) (Active 2 system; Biosemi™, The Netherlands; 10-20 montage). Preprocessing was done using the EEGLAB (version 14.1.1; Delorme & Makeig, 2004) toolbox for MATLAB (version 2017a; MathWorks, Natick, MA). Data were downsampled to 256 Hz, re-referenced to TP8 and filtered using a 1 Hz high pass filter (0.5 Hz transition bandwidth, filter order 1690) and a 45 Hz low pass filter (5 Hz transition bandwidth, filter order 152). Both were zero-phase Hamming windowed sinc FIR filters. Bad channels were automatically detected based on kurtosis measures and rejected after visual confirmation. Artifacts were removed by running an Independent Component Analysis (ICA) to exclude components accounting for motor artifacts, eye blinks, saccades, and bad channels. After ICA, the previously excluded channels were interpolated, using the spherical spline method. Data were segmented into epochs of −100ms to 400ms using a baseline of −100ms to 0ms. These epochs went through an artifact detection algorithm (moving window peal-to-peak threshold at 120 uV). Those subjects with trial rejection percentages above 30% were excluded, which was the case for one subject from the Cystinosis group. No significant differences were found between number of trials included in the analyses per group (Cystinosis group: trial rejection = 7.28%; NT group: trial rejection = 4.93%; *t* = −1.16, *df* = 37.55, *p* = .25).

The definition of the N1 and the MMN windows was based on the typical time of occurrence of the N1 and duration-MMN components, and on visual confirmation that amplitudes were maximal in these intervals. Mean amplitude for the N1 was measured between 80 and 130 ms. The MMN is the difference between deviants and standards and was here measured between 200 and 250 ms (100 to 150 ms post deviance onset). Amplitude measures were taken at FCz, where signal was at its maximum for both responses for both groups. These amplitudes were used for between-groups statistics and correlations. Correlation analyses were computed across groups per component (N1 or MMN) and SOA. All *p-values* (from *post-hoc* tests and correlations) were submitted to Holm-Bonferroni corrections for multiple comparisons (Holm, 1979), using the *p.adjust* of the *stats* package in R (RCoreTeam, 2014).

## Results

Table 1 shows a summary of the included participants’ demographics and performance on the WASI-II. Two-sample independent-means *t* tests were used to test for between-group differences. In cases in which the assumption of the homogeneity of variances was violated, *Welch* corrections were applied to adjust the degrees of freedom. Paired *t* tests were used to test for within-group differences. The IQ statistical analyses revealed significant differences between the groups in all three sub-scales, with individuals diagnosed with Cystinosis showing lower IQ scores on verbal IQ, perceptual reasoning and full scale IQ. The individuals diagnosed with Cystinosis, but not the neurotypical controls, presented significantly lower perceptual reasoning scores, when compared to verbal IQ.

**Table 1.**
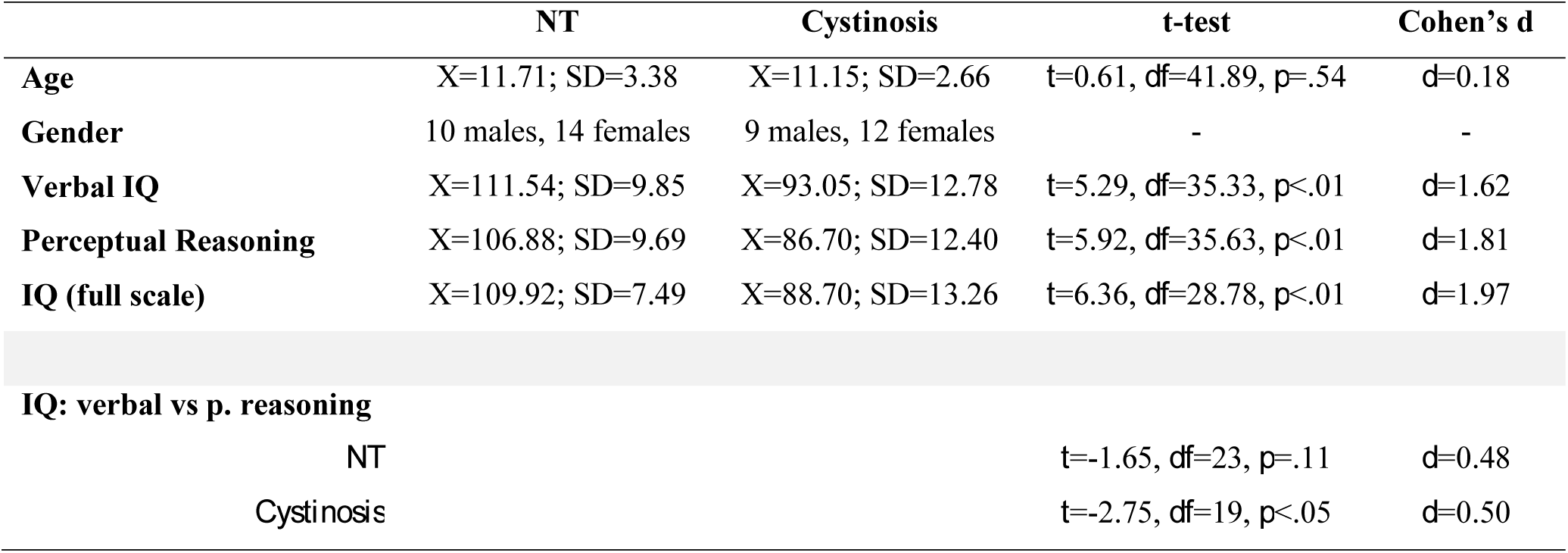
Characterization of the NT and Cystinosis individuals included in the analyses: Age, gender, and IQ

Figure 1 shows the averaged ERPs and topographies for the time windows of interest (N1 and MMN) per SOA and by group.

**Figure1.**
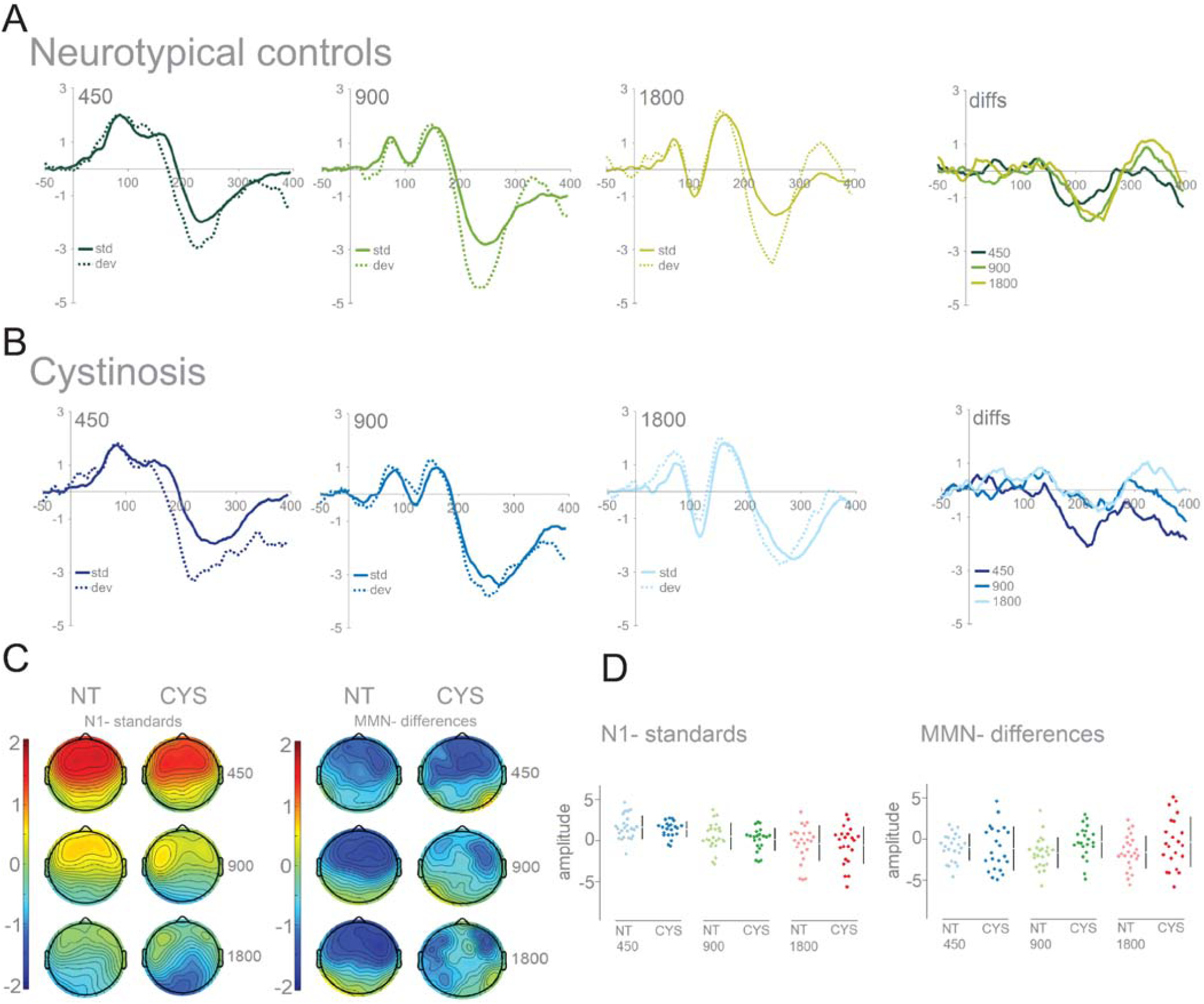
Panel A: Averaged ERPs and topographies per SOA for the NT group at FCz (fourth plot labeled as *diffs* shows MMN, i.e., differences between standards and deviants); Panel B: Averaged ERPs and topographies per SOA for the Cystinosis group at FCz (fourth plot labeled as *diffs* shows MMN, i.e., differences between standards and deviants); Panel C: Topographies for the N1 (standards only) and the MMN time windows, organized from the shorter (450 ms) to the longer (1800 ms) SOA; Panel D: Raw amplitude values are plotted to show distribution. Summary measurements are displayed as gapped lines to the right of each plot: Mean is indicated as a gap in the lines, vertical lines represent standard deviation error bars.

Mixed-effects models were implemented to analyze the EEG data, using the *lmer* function in the lme4 package (Bates, Mächler, Bolker, & Walker, 2014) in R (Version 3.1.2, (RCoreTeam, 2014)). The models were run separately for the N1 and the MMN time windows. Mean amplitude at FCz was the numeric dependent variable. For the N1, only standard amplitudes were considered. For the MMN, mean amplitude referred to amplitude of the difference between standards and deviants. Group (NT = −0.5, Cystinosis = 0.5) was a contrast-coded fixed factor, and SOA was a numeric fixed factor. Subject and SOA were random factors. While the inclusion of SOA as a fixed effect measures the overall effect of SOA on amplitude, its inclusion as a random effect aims to account for SOA variance and the impact of that variance on the fixed effects and on the fit of the model. Models were fit using the maximum likelihood criterion. *P* values were estimated using Satterthwaite approximations (Satterthwaite, 1946).

As expected, in the N1 time window there was a significant effect of SOA, with both 900 ms (*ß* = −1.20, SE = 0.07, *p* < .001) and 1800 ms (*ß* = −2.02, SE = 0.07, *p* < .001) conditions eliciting more negative responses than the shortest SOA (450 ms). No effect of group or interaction between group and SOA was found. No significant correlations were found with IQ.

In the MMN time window, there was a significant interaction between group and SOA, with the difference between 450ms and 900ms (*ß* = 1.86, SE = 0.18, *p* < .01) and between 450ms and 1800ms (*ß* = 1.30, SE = 0.18, *p* < .01) being larger for the individuals diagnosed with Cystinosis than for the neurotypical controls. *Post-hoc* analyses revealed that this was due to significantly decreased responses for the 900ms and 1800ms SOAs in individuals diagnosed with Cystinosis, when compared to the neurotypical controls (*t* = −7.54, *df* = 567.04, *p* < .01); (*t* = −4.54, *df* = 481.59, *p* < .01). As for N1, no significant correlations were found between MMN and IQ.

## Discussion

We used high-density EEG and a passive oddball paradigm to characterize early auditory sensory processing and sensory memory in a sample of children and adolescents with Cystinosis. Additionally, we measured the associations between auditory brain function and verbal and non-verbal IQ.

No differences were found between the groups in the auditory N1, suggesting that sensory transmission through the auditory system is largely intact in individuals with Cystinosis. This is in accordance with preliminary data from our lab showing maintained auditory processing in the context of a multisensory task in a modest sample of individuals with Cystinosis (Andrade et al., 2016). Though enhanced auditory potentials have been described for this population (Čeponiene et al., 2008), such differences were observed in a later, functionally distinct component (P2), in a sample with a significantly wider range of ages, and during a task focused on spatial selective attention. As can be seen in Figure 1, our data do not seem to support the presence of an enhanced P2 in the current sample. Further, here, N1 was shown to modulate as a function of SOA in both Cystinosis and neurotypical control groups. This has been consistently described in the literature for the neurotypical population (Teder, Alho, Reinikainen, & Näätänen, 1993) and is often explained by one of two (non-exclusive) alternatives: habituation (Özesmi, Dolu, Süer, Gölgelio, & Aşçioglu, 2000; Thompson & Spences, 1966) or refraction (Budd, Barry, Gordon, Rennie, & Michie, 1998; Tremblay, Billings, & Rohila, 2004). Our findings therefore indicate highly typical auditory sensory response properties in Cystinosis.

In the MMN time window, significantly decreased responses were found in the Cystinosis group for the two longer SOAs, whereas a robust MMN was elicited for the shortest SOA. Previous work from our lab using this same MMN paradigm showed that increasing SOA similarly led to diminution of the MMN in Rett Syndrome (Brima et al., 2019), such that the MMN was no longer detectable at the longer SOAs. This was taken to index weakened maintenance of the memory trace in Rett. That differences between the groups were here, likewise, only observed for the longer SOAs, suggests that Cystinosis (at least during childhood and adolescence) might be characterized by deficits in the maintenance of short term auditory sensory memory (Bartha-Doering, Deuster, Giordano, am Zehnhoff-Dinnesen, & Dobel, 2015).

An impairment in auditory sensory memory (a preattentive memory system that allows an individual to retain traces of sensory information after the termination of the original stimulus (Cowan, 1999)) could impact subsequent processing in working memory (Bonetti et al., 2018), a conscious cognitive system responsible for the temporary holding, processing, and manipulation of information (Baddeley, 1992). Indeed, despite being somewhat separate processes with unique characteristics, auditory sensory memory and working memory seem to be associated in neurotypical controls–with those individuals who show better performance in working memory tasks, presenting enhanced MMN responses (Bonetti et al., 2018)–and in clinical populations, with impaired auditory MMN being associated with deficits in working memory (Ahveninen et al., 1999; Javitt, Doneshka, Grochowski, & Ritter, 1995). Furthermore, they have been suggested to share neural bases (Pasternak & Greenlee, 2005). The deficit in auditory sensory memory reported here would thus be seemingly at odds with previous evidence of an enhanced auditory working memory in a modest sample of individuals with Cystinosis (Nichols, Press, Schneider, & Trauner, 1990). In a memory for sentences task (which asks the individual to recall sentences of increasing length and complexity), children and adolescents diagnosed with Cystinosis performed better than in other subscales of the Stanford-Binet, which was argued as a potential compensatory mechanism for their poorer visual memory (Nichols et al., 1990). Nevertheless, no neurotypical controls were assessed and, therefore, though those with Cystinosis showed higher scores in the memory for sentences task than in the additional tasks, this finding does not allow one to draw conclusions about the typicality of such scores. And, indeed, in a study comparing children diagnosed with Cystinosis with their neurotypical peers, working memory, as assessed by a parent-completed questionnaire, appeared to be a problematic area in those with Cystinosis (Ballantyne et al., 2013).

Impaired auditory sensory memory could, ultimately, hamper language acquisition and processing (Čeponiene et al., 1999). Considering the average verbal performance in the individuals tested here and, generally, in Cystinosis, one might nevertheless argue that, in this population, despite the presence of early auditory sensory memory differences, the system appears to be resilient and to compensate for those differences at a later stage of processing. Future work addressing other aspects of sensory and working memory will be needed to better understand the underlying bases of the differences described here, and their implication for higher order processing.

Lastly, despite an average verbal IQ, individuals with Cystinosis presented low average non-verbal (perceptual) and full-scale IQ scores. Other studies have reported significantly lower IQs in this population, when compared to neurotypical controls (Aly et al., 2014; Ulmer et al., 2009). Although the exact cause of the cognitive deficits observed in Cystinosis is unknown, early cystine accumulation might be particularly detrimental to brain myelination through in utero damage to pre-oligodendroglial cells, which are susceptible to the type of oxidative stress resulting from the metabolic impairment associated with CTNS mutations (Trauner et al., 2007). Myelination atypicalities could subsequently hinder the development of cortical projections crucial for complex cognitive processes (Trauner et al., 2007). The expected discrepancy between verbal and non-verbal indices (Frankel & Trauner, 2019; Spilkin et al., 2007; Ulmer et al., 2009) was also observed in the current study. Such a discrepancy could be explained by abnormal white mater microstructure in visual-related areas: In a diffusion tensor imaging study, children diagnosed with Cystinosis presented decreased fractional anisotropy and increased mean diffusivity in the dorsal visual pathway (Bava et al., 2010). Of note, however, IQ scores did not correlate significantly with our neural measures of interest (the N1 and MMN), suggesting that both basic auditory processing and sensory memory are not strongly associated with verbal and non-verbal abilities, at least as measured here. A dissociation between MMN and IQ was has been previously shown in a study comparing neural responses of children with intellectual disability, developmental dysphasia, and neurotypical controls (Holopainen, Korpilahti, Juottonen, Lang, & Sillanpää, 1998).

Several limitations to the present study should be addressed in future research. First, despite the substantial size of our sample considering the rare nature of Cystinosis, larger numbers would allow for more detailed analyses, particularly those looking at associations between neural, cognitive and behavioral outcomes. Furthermore, it will be important to characterize the developmental trajectory of auditory sensory memory and the potential impact of continued treatment-associated factors (dialysis, number of transplants) and/or of cystine accumulation in the brain, with a larger sample that includes adults and younger children. While identification of differences in sensory memory provide a potential biomarker of the effects of cysteamine on brain function to serve as secondary outcome measure for clinical trials, it will be critical to determine whether the deficit varies with other clinically significant symptoms. Furthermore, it will be interesting to determine if similar deficits are seen in unaffected carriers of the mutation, as shown for visual-spatial difficulties (Sathappan & Trauner, 2019), or if they are specific to the effects of cysteamine accumulation. As alluded to above, future work will need to be done to understand the implications of the auditory sensory memory deficit described here. For example, one would ideally have other auditory sensory and working memory measures supportive of these difficulties.

In summary, this study provides the first neural evidence of auditory sensory memory differences in children and adolescents diagnosed with Cystinosis, which has the potential to serve as a biomarker of the effects of treatment and of cysteamine on brain function.

## Author Contributions

JJF and SM conceived the study and designed the original experiment. AAF and DJH collected and analyzed the data. AAF wrote the first draft of the manuscript. SM and JJF provided editorial input to AAF on the subsequent drafts.

## Acknowledgements

We wish to thank Drs. Juliana Bates and Katherine Behar, who performed the clinical assessments, and Elise Taverna for her help with data collection. We extend our most sincere gratitude to the participants and their families for their interest, their involvement, and their time.

This work was supported by a grant from the Cystinosis Research Network and a Eunice Kennedy Shriver National Institute of Child Health and Human Development U54 Grant (HD090260) to the Human Clinical Phenotyping Core of the Rose F. Kennedy Intellectual and Developmental Disabilities Research center.

## References

Ahveninen, J., Jääskeläinen, I. P., Pekkonen, E., Hallberg, A., Hietanen, M., Mäkelä, R., … Sillanaukee, P. (1999). Suppression of mismatch negativity by backward masking predicts impaired working□memory performance in alcoholics. Alcoholism: Clinical and Experimental Research, 23(9), 1507–1514.

Aly, R., Makar, S., El Bakri, A., & Soliman, N. A. (2014). Neurocognitive functions and behavioral profiles in children with nephropathic cystinosis. Saudi Journal of Kidney Diseases and Transplantation, 25(6), 1224–1231.

Andrade, G. N., Molholm, S., Pal, A., Kaskel, F., Walkley, S. U., & Foxe, J. J. (2016). Multisensory processing in isorders: A behavioral and high-density electrophysiology investigation in Niemann-Pick type C and cystinosis. Paper presented at the WORLDSymposium, San Diego, CA.

Baddeley, A. (1992). Working memory. Science, 255, 556–559.

Ballantyne, A., Scarvie, K. M., & Trauner, D. A. (1997). Academic achievement in individuals with infantile nephropathic cystinosis. American Journal of Medical Genetics (Neuropsychiatric Genetics), 74, 157–161.

Ballantyne, A., Spilkin, A. M., & Trauner, D. A. (2013). Executive function in nephropathic cystinosis. Cognitive and Behavioral Neurology, 26(1), 14–22.

Bartha-Doering, L., Deuster, D., Giordano, V., am Zehnhoff-Dinnesen, A., & Dobel, C. (2015). A systematic review of the mismatch negativity as an index for auditory sensory memory: From basic research to clinical and developmental perspectives. Psychophysiology, 52, 1115–1130.

Bates, D., Mächler, M., Bolker, B. M., & Walker, S. C. (2014). Ime4: Linear mixed-effects models using Eigen and S4 (R package Version 1.1-7) [Computer software]. Retrieved from http://CRAN.R-project.org/package=lme4.

Bava, S., Theilmann, R. J., Sach, M., May, S. J., Frank, L. R., Hesselink, J. R., … Trauner, D. A. (2010). Developmental changes in cerebral white matter microstructure in a disorder of lysosomal storage. Cortex, 46, 206–216.

Besouw, M. T. P., Hulstijn-Dirkmaat, G. M., van der Rijken, R. E. A., Cornelissen, E. A. M., van Dael, C. M., Vande Walle, J., … Levtchenko, E. N. (2010). Neurocognitive functioning in school-aged cystinosis patients. Journal of Inherited Metabolic Disease, 33, 787–793.

Bonetti, L., Haumann, N. T., Brattico, E., Kliuchko, M., Vuust, P., Särkämö, T., & Näätänen, R. (2018). Auditory sensory memory and working memory skills: Association between frontal MMN and performance scores. Brain Research, 1700, 86–98.

Brima, T., Molholm, S., Molloy, C. J., Sysoeva, O. V., Nicholas, E., Djukic, A., … Foxe, J. J. (2019). Auditory sensory memory span for duration is severely curtailed in females with Rett syndrome. Translational Psychiatry, 9(1), 130.

Budd, T. W., Barry, R. J., Gordon, E., Rennie, C., & Michie, P. T. (1998). Decrement of the N1 auditory event-related potential with stimulus repetition: habituation vs. refractoriness. International Journal of Psychophysiology, 31, 51–68.

Čeponiene, R., Hukki, J., Cheour, M., Haapanen, M. L., Ranta, R., & Näätänen, R. (1999). Cortical auditory dysfunction in children with oral clefts: relation with cleft type. Clinical Neurophysiology, 110(11), 1921–1926.

Čeponiene, R., Trauner, D., Kinnear, M., Spilkin, A., Jong, K., Girard, H., & Townsend, J. (2008). Perception, attention, and target detection in auditory and visual modalities in cystinosis. Paper presented at the First International Cystinosis Research Symposium Irvine, California.

Cochat, P., Drachman, R., Gagnadoux, M. F., Pariente, D., & Broyer, M. (1986). Cerebral atrophy and nephropathic Cystinosis. Archives of Disease in Childhood, 61(4), 401–403.

Cowan, N. (1999). An embedded-processes model of working memory. In A. Miyake & P. Shah (Eds.), Models of Working Memory: Mechanisms of Active Maintenace and Executive Control (pp. 62–101). Cambridge, UK: Cambridge University Press.

Ehrich, J. H. H., Stoeppler, L., Offner, G., & Brodehl, J. (1979). Evidence for cerebral involvement in nephropathic Cystinosis. Neuropadiatrie, 10(2), 128–137.

Elmonem, M. A., Veys, K. R., Soliman, N. A., van Dyck, M., van den Heuvel, L., & Levtchenko, E. (2016). Cystinosis: A review. Orphanet Journal of Rare Diseases, 11(47), 1–12.

Ethier, A., Lippé, S., Mérouani, A., Lassonde, M., & Saint-Amour, D. (2012). Reversible visual evoked potential abnormalities in uremic children. Pediatric Neurology, 46, 390–392.

Fink, J. K., Brouwers, P., Barton, N., Malekzadeh, M. H., Sato, S., Hill, S., … Gahl, W. A. (1989). Neurologic complications in long-standing neohropathic cystinosis. Archives of Neurology, 46, 543–548.

Frankel, A. M., & Trauner, D. A. (2019). Visual and verbal learning and memory in cystinosis. Brain and Cognition, 135, 103578.

Gahl, W. A. (1986). Cystinosis coming of age. Advances in Pediatrics, 33, 95–126.

Gahl, W. A., Bashan, N., Tietze, F., Bernardini, I., & Schulman, J. D. (1982). Cystine transport is defective in siolated leukocyte lysosomes from patients with cystinosis. Science, 217, 1263–1265.

Gahl, W. A., & Kaiser-Kupfer, M. I. (1987). Complications of nephropathic cystinosis after renal failure. Pediatric Nephrology, 1, 260–268.

Gahl, W. A., Thoene, J. G., & Schneider, J. A. (2009). Cystinosis. The New England Journal of Medicine, 347(2), 111–121.

Giard, M. H., Perrin, F., Echallier, J. F., Thevenet, M., Froment, J. C., & Pernier, J. (1994). Dissociation of temporal and frontal components in the human auditory N1 wave: A scalp current density and dipole model analysis. Electroencephalography and Clinical Neurophysiology, 92, 238–253.

Holm, S. (1979). A Simple Sequentially Rejective Multiple Test Procedure. Scandinavian Journal of Statistics, 6(2), 65–70.

Holopainen, I. E., Korpilahti, P., Juottonen, K., Lang, H., & Sillanpää, M. (1998). Abnormal frequency mismatch negativity in mentally retarded children and in children with developmental dysphasia. Journal of Child Neurology, 13(4), 178–183.

Ivanova, E., De Leo, M. G., De Matteis, M. A., & Levtchenko, E. (2014). Cystinosis: Clinical presentation, pathogenesis and treatment. Pediatric Endocrinology Reviews, 11(Suppplement 1), 387–394.

Javitt, D. C., Doneshka, P., Grochowski, S., & Ritter, W. (1995). Impaired mismatch negativity generation reflects widespread dysfunction of working memory in schizophrenia. Archives of General Psychiatry, 52(7), 550–558.

Jonas, A. J., Greene, A. A., Smith, M. L., & Schneider, J. A. (1982). Cystine accumulation and loss in normal, heterozygous, and cystinotic fibroblasts. Proceedings of the National Academy of Sciences of the United States of America, 79, 4442–4445.

Leavitt, V. M., Molholm, S., Ritter, W., Shpaner, M., & Foxe, J. J. (2007). Auditory processing in schizophrenia during the middle latency period (10–50 ms): high-density electrical mapping and source analysis reveal subcortical antecedents to early cortical deficits. Journal of Psychiatry & Neuroscience, 32(5), 339–353.

Levine, S., & Paparo, G. (1982). Brain lesions in a case of Cystinosis Acta Neuropathologica, 57, 217–220.

Levtchenko, E., van den Heuvel, L., Emma, F., & Antignac, C. (2014). Clinical utility gene card for: Cystinosis. European Journal of Human Genetics, 22, e1–e3.

Müller, M., Baumeier, A., Ringelstein, E. B., & Husstedt, I. W. (2008). Long-term tracking of neurological complications of encephalopathy and myopathy in a patient with nephropathic cystinosis: a case report and review of the literature. Journal of Medical Case Reports, 2, 235.

Näätänen, R., & Alho, K. (1995). Mismatch negativity-A unique measure of sensory processing in audition. International Journal of Neuroscience, 80(1-4), 317–337.

Näätänen, R., & Picton, T. (1987). The N1 wave of the human electric and magnetic response to sound: A review and an analysis of the component structure. Psychophysiology, 24(4), 375–425.

Näätänen, R., & Winkler, I. (1999). The concept of auditory stimulus representation in cognitive neuroscience. Psychological Bulletin, 125(6), 826–859.

Nichols, S. L., Press, G. A., Schneider, J. A., & Trauner, D. A. (1990). Cortical atrophy and cognitive performance in infantile nephropathic cystinosis. Pediatric Neurology, 6(6), 379–381.

Niemiec, S., Ballantyne, A., & Trauner, D. A. (2012). Cognition in nephropathic cystinosis: Pattern of expression in heterozygous carriers. American Journal of Medical Genetics Part A, 158(8), 1902–1908.

Özesmi, Ç., Dolu, N., Süer, C., Gölgelio, A., & Asçioglu, M. (2000). Habituation of the auditory evoked potential in a short interstimulus interval paradigm. International Journal of Neuroscience, 105, 87–95.

Pasternak, T., & Greenlee, M. W. (2005). Working memory in primate sensory systems. Nature Reviews Neuroscience, 6, 97–107.

Rao, K. I., Hesselink, J. R., & Trauner, D. A. (2015). Chiari I malformation in nephropathic cystinosis. The Journal of Pediatrics, 167, 1126–1129.

R Core Team. (2014). R: A language and environment for statistical computing (Version 3.3.2). Vienna, Austria: R Foundation for Statistical Computing.

Ritter, W., Deacon, D., Gomes, H., Javitt, D. C., & Vaughan, J. H. (1995). The mismatch negativity of event-related potentials as a probe of transient auditory memory: A review. Ear and Hearing, 16(1), 52–67.

Ritter, W., Sussman, E., Molholm, S., & Foxe, J. J. (2002). Memory reactivation or reinstatement and the mismatch negativity. Psychophysiology, 39, 158–165.

Sathappan, A., & Trauner, D. A. (2019). Hierarchical processing of visual stimuli in nephropathic Cystinosis. Journal of Inherited Metabolic Disease, 42(3), 545–552.

Satterthwaite, F. E. (1946). An approximate distribution of estimates of variance components. Biometrics Bulletin, 2, 110–114.

Scarvie, K. M., Ballantyne, A., & Trauner, D. A. (1996). Visuomotor performance in children with infantile nephropathic Cystinosis. Perceptual and Motor Skills, 82, 67–75.

Schneider, J. A., Katz, B., & Melles, R. B. (1990). Update on nephropathic Cystinosis. Pediatric Nephrology, 4(6), 645–653.

Shotelersuk, V., Larson, D., Anikster, Y., McDowell, G., Lemons, R., Bernardini, I., … Gahl, W. A. (1998). CTNS mutations in an American-based population of Cystinosis patients. American Journal of Human Genetics, 63, 1352–1362.

Spilkin, A. M., Ballantyne, A. O., Babchuck, L. R., & Trauner, D. A. (2007). Non□verbal deficits in young children with a genetic metabolic disorder: WPPSI□III performance in Cystinosis. American Journal of Medical Genetics Part B: Neuropsychiatric Genetics, 144(4), 444–447.

Teder, W., Alho, K., Reinikainen, K., & Näätänen, R. (1993). Interstimulus interval and the selective-attention effect on auditory ERP: “N1 enhancement” versus processing negativity. Psychophysiology, 30, 71–81.

Theodoropoulos, D. S., Krasnewich, D., Kaiser-Kupfer, M. I., & Gahl, W. A. (1993). Classic nephropathic Cystinosis as an adult disease. The Journal of the American Medical Association, 270(18), 2200–2204.

Thompson, R. F., & Spences, W. A. (1966). Habituation: A model phenomenon for the study of neuronal substrates of behavior. Psychological Review, 73(1), 16–43.

Town, M., Jean, G., Cherqui, S., Attard, M., Forestier, L., Whitmore, S. A., … Antignac, C. (1998). A novel gene encoding an integral membrane protein is mutated in nephropathic cystinosis. Nature Genetics, 18, 319–324.

Trauner, D. A., Chase, C., Scheller, J., Katz, B., & Schneider, J. A. (1988). Neurologic and cognitive deficits in children with Cystinosis. The Journal of Pediatrics, 112(6), 912–914.

Trauner, D. A., Spilkin, A. M., Williams, J., & Babchuck, L. (2007). Specific cognitive deficits in young children with cystinosis: Evidence for an early effect of the cystinosin gene on neural function. The Journal of Pediatrics, 151(2), 192–196.

Trauner, D. A., Williams, J., Ballantyne, A., Spilkin, A. M., Crowhurst, J., & Hesselink, J. R. (2010). Neurological impairment in nephropathic cystinosis: motor coordination deficits. Pediatric Nephrology, 25, 2061–2066.

Tremblay, K. L., Billings, C., & Rohila, N. (2004). Speech evoked cortical potentials: Effects of age and stimulus presentation rate. Journal of the American Academy of Audiology, 15, 226–237.

Ulmer, F. F., Landolt, M. A., Ha Vinh, R., Huisman, T. A. G. M., Neuhaus, T. J., Latal, B., & Laube, G. F. (2009). Intellectual and motor performance, quality of life and psychosocial adjustment in children with cystinosis. Pediatric Nephrology, 24, 1371–1378.

van Rijssel, A. E., Knuijt, S., Veys, K., Levtchenko, E. N., & Janssen, M. C. H. (2019). Swallowing dysfunction in patients with nephropathic cystinosis. Molecular Genetics and Metabolism, 126, 413–415.

Viltz, L., & Trauner, D. A. (2013). Effect of age at treatment on cognitive performance in patients with Cystinosis. The Journal of Pediatrics, 163, 489–492.

Vogel, D. G., Malekzadeh, M. H., Cornford, M. E., Schneider, J. A., Shields, W. D., & Vinters, H. V. (1990). Central nervous system involvement in nephropathic Cystinosis. Journal of Neuropathology and Experimental Neurology, 49(6), 591–599.

